# Increased H-Bond Stability Relates to Altered ε-Cleavage Efficiency and Aβ Levels in the I45T Familial Alzheimer’s Disease Mutant of APP

**DOI:** 10.1101/372698

**Authors:** Alexander Götz, Philipp Högel, Mara Silber, Iro Chaitoglou, Burkhard Luy, Claudia Muhle-Goll, Christina Scharnagl, Dieter Langosch

## Abstract

Cleavage of the amyloid precursor protein’s (APP) transmembrane domain (TMD) by γ-secretase is a crucial step in the aetiology of Alzheimer’s Disease (AD). Mutations in the APP TMD alter cleavage and lead to familial forms of AD (FAD). The majority of FAD mutations shift the preference of initial cleavage from ε49 to ε48, thus raising the AD-related Aβ42/Aβ40 ratio. The I45T mutation is among the few FAD mutations that do not alter ε-site preference, while it dramatically reduces the efficiency of ε-cleavage. Here, we investigate the impact of the I45T mutation on the backbone dynamics of the substrate TMD. Amide exchange experiments and molecular dynamics simulations in solvent and a lipid bilayer reveal an increased stability of amide hydrogen bonds at the ζ-and γ-cleavage sites. Stiffening of the H-bond network is caused by an additional H-bond between the T45 side chain and the TMD backbone, which alters dynamics within the cleavage domain. In particular, the increased H-bond stability inhibits an upward movement of the ε-sites in the I45T mutant. Thus, an altered presentation of ε-sites to the active site of γ-secretase as a consequence of restricted local flexibility provides a rationale for reduced ε-cleavage efficiency of the I45T mutant.

## Introduction

The intramembrane aspartyl protease γ-secretase cleaves the transmembrane domains (TMD) of ~90 bitopic type I transmembrane proteins^1,2^. Due to its involvement in Alzheimer’s disease (AD), the amyloid precursor protein (APP) is the most extensively studied γ-secretase substrate^3,4^. Prior to cleavage by γ-secretase, APP’s ectodomain is shed by β-secretase, thus generating the C99 fragment. Subsequent cleavage of C99 by γ-secretase results in the release of the APP intracellular domain (AICD) and β-amyloid (Aβ) peptides of various lengths^5^. Such step-wise cleavage starts at one of two ε-sites located at the cytosolic border of the C99 TMD between either residues L49 and V50 (ε49) or T48 and L49 (ε48). Cleavage gradually proceeds towards the N-terminus, releasing fragments of three or four residues and leaving Aβ peptides of different lengths^6–11^. Aβ40, the most abundant Aβ peptide produced from wild-type (WT) C99, is generated by cleavage along the ε49-ζ46-γ43-γ40 pathway. A smaller amount of C99 is processed via cleavage at ε48 and ζ45, leading to Aβ42 and Aβ38 peptides. Of these species, Aβ42 is the most aggregation-prone and forms neurotoxic oligomers and plaques^12,13^. The accumulation of plagues consisting of such cell-toxic Aβ peptides in the brain is a central hallmark of AD^14^. In familial forms of AD (FAD), an increased Aβ42/Aβ40 ratio correlates with early onset and fast progression of AD and results from point mutations in C99 or presenilin, the catalytic subunit of γ-secretase^1,15–21^. In most mutants, alterations in Aβ42/Aβ40 ratios are linked to shifting the preferential initial cleavage site from ε49 to ε48. However, switching pathways after ε-cleavage is also seen^8,9,11,22^. The I45T mutation is one case where an increased Aβ42/Aβ40 ratio results from pathway switching^23^. In addition, I45T dramatically reduces cleavage efficiency at both ε-sites^20,21,23^.

Several studies have indicated that the C-terminal part of the C99 TMD helix (TM-C), which comprises the cleavage sites, is less flexible than the N-terminal region (TM-N) ^24–29^. As cleavage of the peptide backbone requires that presenilin’s catalytic aspartates are able to access the scissile bond^30^, a mutationally altered structural stability of the C99 TMD helix at the ε-sites seemed to be an obvious explanation for altered ε-cleavage^28,31,32^. In addition, FAD mutations in TM-C have been shown to alter hydrogen bond (H-bond) stability in a segment upstream of the ε-cleavage sites which harbours the γ sites^29,33^. Those alterations relate to changed shape fluctuations of the TMD helix, which are controlled by the previously established G_37_G_38_ hinge^26,29,34,35^ and by a flexible site around T_43_-I_45_^29^.

Previously, it was shown that Thr residues in the WT C99 TMD (T43 and T48) rigidify the helix by forming H-bonds between their side chains and the main chain^26,35^. Mutations of T43 and T48 to hydrophobic residues (e.g., Val or Ile) affect ε-site preference and efficiency^15,21,36^. Since hydrogen bond networks determine the mechanical and thermodynamic properties of proteins, their alteration in FAD mutants of the APP TMD has been proposed to perturb the fine-tuned interplay of the conformational dynamics of the enzyme and substrate^29^, which is a key factor for substrate recognition and subsequent relaxation steps^37,38^.

Here, we combined amide exchange experiments with atomistic simulations in order to compare the conformational dynamics of WT and I45T mutant TMD helices. Our data confirm a more flexible TM-N compared to the very rigid TM-C. This agrees with previous results^26,29,39–41^ but contradicts a novel interpretation of amide exchange experiments in a very recent study^42^. The I45T mutation stabilizes H-bonds in TM-C mainly around the γ-sites. As expected, this stabilization is related to a newly formed H-bond between the T45 side chain and the backbone carbonyl oxygen of I41. Around the ε-sites, neither H-bond stability nor local flexibility was affected. Exhaustive sampling of the conformational space allowed us to correlate the impact of the I45T mutation on the H-bond network with altered bending and twisting motions around a flexible hinge located one turn upstream of the ε-sites. These motions may control the vertical position of the ε-cleavage sites within presenilin and thus provide a mechanistic interpretation for changed ε-cleavage efficiency but unaffected ε-site preference, in the I45T mutant.

## Results

In order to provide a rationale for the altered γ-secretase cleavage of the C99 I45T FAD mutant, we probed the structure and flexibility of its TMD in comparison to the WT TMD. For this purpose, a 30 amino acid long fragment of C99 was used, which covers residues S26 - K55 (A26-55). Similar C99 TMD peptides were previously shown to be good substrates for γ-secretase^31,43^. Circular dichroism (CD) spectroscopy, amide exchange experiments performed by mass spectrometry (MS) as well as NMR spectroscopy, and molecular dynamics (MD) simulations in the μs range were combined to study the TMD helix dynamics. The polar environment of the catalytic cleft of presenilin was mimicked by aqueous 80% 2,2,2-trifluoroethanol (TFE)^25–27,44,45^. This solvent mimics the polarity in the solvated interior of proteins^29,44,46,47^. Simulations were performed in the same solvent and in a 1-palmitoyl-2-oleoylphosphatidylcholine (POPC) bilayer. The comparison of TMD dynamics in both environments extends our previous *in silico* modelling study of the backbone dynamics of the C99 WT TMD and its FAD mutations^29^. Performing amide exchange in membranes was not practical as the bilayer shields the TMD helix core from exchange^48,49^.

### The I45T FAD mutation locally reduces helix flexibility

CD spectra indicate that the peptides are strongly helical in 80% TFE (Supplementary Fig. S2). Normalized spectra reveal that WT and I45T share equivalent helicity. To assess the TM-helices’ conformational flexibility, intrahelical amide H-bond stabilities were probed by amide deuterium hydrogen exchange (DHX) in 80% TFE. For DHX, exhaustively (>98%) deuterated A26-55 peptides were diluted to a concentration of 5 μM into a protonated solvent, which leaves them in the monomeric state^27^. Exchange of amide deuterons to protons was recorded over time at pH 5.0 and 20°C. Gradual shifting of monomodal isotopic envelopes towards lower m/z values was detected (data not shown). This is characteristic of EX2 kinetics with uncorrelated exchange of individual deuterons upon local unfolding^50,51^. A qualitative comparison of the overall DHX kinetics revealed nearly identical exchange kinetics for the rapidly exchanging deuterons (0-3 h) of WT and I45T (Fig. 1a, inset). With increasing incubation time, I45T showed slower exchange than WT, with a difference of ~ 1 D after 6 h. At later time points, I45T showed significantly faster exchange, resulting in the curves crossing at ~16 h. MD simulations reproduced the overall DHX kinetics well (Supplementary Fig. S3a,b).

**Figure 1.**
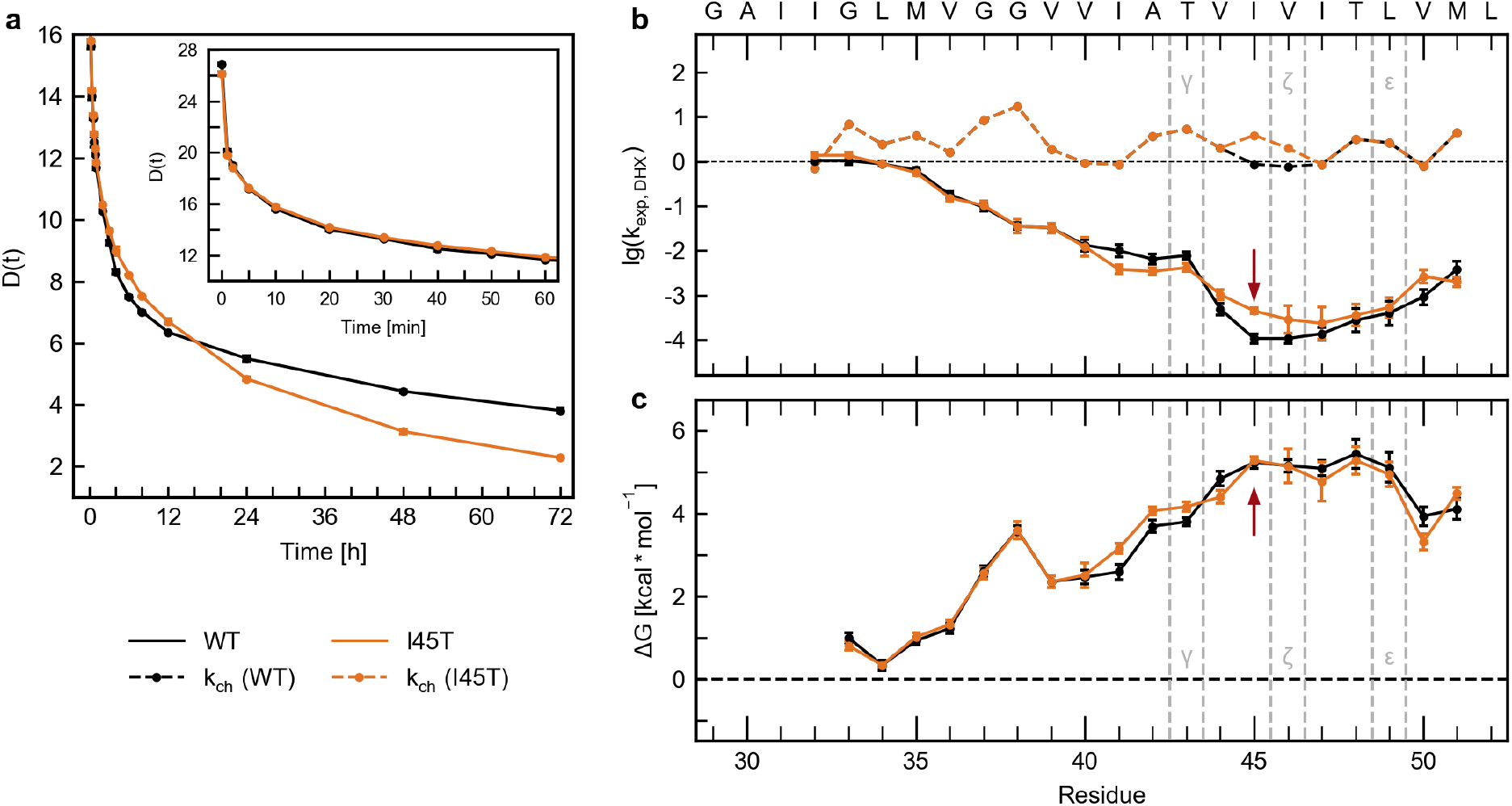
Deuterium-hydrogen exchange kinetics and H-bond stabilities determined by ESI-TOF MS. (**a**) Overall amide DHX kinetics of A26-55. The number of remaining deuterons plotted as a function of time for the initial 60 min (inset) and the complete 72 h of incubation (mean values, n ≥ 3, SEM smaller than the symbols). (**b**) DHX rate constants k_exp,DHX_ [1/min] of individual amide deuterons from ETD measurements (n ≥ 3, mean values ± SE). The dashed lines correspond to the calculated chemical amide exchange rate constants k_ch_ [1/min]. (c) Strength (ΔG) of intrahelical amide H-bonds based on calculated exchange rate constants. Calculation of ΔG was not possible for residues G29-I32 at the N-terminus where k_ch_ exceeded the experimental exchange rate constants. Error bars correspond to standard confidence intervals calculated from the standard errors of k_exp,DHX_.

To map the impact of the I45T mutation on local helix flexibility, site-specific amide DHX rate constants (k_exp,DHX_) were measured. After different periods of DHX in solution (0.1 min to 7 d at pH 5.0), we determined the deuteron contents of the c- and z-fragments resulting from electron transfer dissociation (ETD) in the gas phase. Residue-specific k_exp,DHX_ rate constants were calculated based on the respective mass shifts over time. As ETD of the original A26-55 peptide was insufficient, we exchanged S26 and N27 to lysine (termed A28-55 in the following). To exclude major consequences of this sequence alteration on TMD dynamics, we compared the overall DHX of A28-55 with the DHX of A26-55. Excellent agreement was found for intermediate and long incubation times while some deviations were observed for rapidly exchanging deuterons (Supplementary Fig. S3c).

The k_exp,DHX_ rate profile of A28-55 reveals a rapidly exchanging TM-N, which is followed by a rather stable TM-C that harbours the γ-secretase cleavage sites (Fig. 1b). The k_exp,DHX_ values are below the respective chemical amide exchange rate constants k_ch_ from G33 through M51, which indicates the participation of these residues in the secondary structure, *viz*. helix formation. To validate our determination of ETD-based exchange rate constants, we reconstructed the overall DHX kinetics from k_exp,DHX_ rate constants of A28-55 WT. The reconstructed kinetics show very good agreement with directly measured overall kinetics, except for very slowly exchanging deuterons that exchanged somewhat too rapidly in the reconstructed kinetics (Supplementary Fig. S3d). Reconstructed overall exchange kinetics also reproduced the crossing of the overall WT and I45T DHX curves (Supplementary Fig. S3e). A faster exchange of TM-N over TM-C was confirmed by solution NMR measurements (Supplementary Fig. S4), using H_α_/HN (TOCSY) or HN/N (^1^H^15^N-HSQC) cross peak measurements on WT A26-55 with ^15^N/^13^C labels at positions G29, G33, G37, G38, I41, V44, M51, and L52.

MS-ETD measurements were then employed to compare WT and I45T TMDs. For the I45T mutant, lower k_exp,DHX_ rate constants were observed from I41 to T43, while an increased rate constant was found for T45. This is in perfect agreement with the overall exchange kinetics (Fig. 1a), assuming that I41-T43 contributes to the fast population of amides (exchanging <16 h), and that T45 contributes to the slow population (>16 h).

The rate constants k_exp,DHX_ contain information on the stability of the respective amide H-bonds, the local concentration of the exchange catalyst as well as the chemical exchange rate kch. kch, in turn, depends on (1) the pH and (2) side-chain chemistry, i.e., primary structure. The effects of primary structure are taken into account by converting the rate constants into the free energy change of amide H-bond formation (ΔG), which is a measure of H-bond stability, by the Linderstrøm-Lang theory^52^ (Fig. 1c). We note that derived ΔG values represent upper estimates of the true values^53^. The distribution of ΔG values across the sequence confirms the existence of a highly flexible, Gly-rich TM-N (ΔG < 2 kcal/mol), followed by a rigid TM-C (ΔG ~5 kcal/mol). The I45T mutation induces a slight increase in stability from I41 to T43 (ΔΔG ≈0.3 kcal/mol). No differences in ΔG are seen around position 45, which indicates that the difference of k_exp,DHX_ at position 45 results from a difference in k_ch_ between Thr and Ile. Further, no difference is seen at the ε- or ζ-cleavage sites.

In a different approach to assess the conformational flexibility of the C99 TMD, Cao et al.^42^ recently reported site-specific isotope fractionation factors Φ obtained from the ratio of NMR-derived hydrogen deuterium exchange (HDX) and DHX amide exchange rate constants that were recorded in lyso-myristoylphosphatidylglycerol (LMPG) micelles^42^. Generally, the Φ value reports the isotopic preference in an H-bond and it is often assumed that the equilibrium enrichment of deuterium at the amide (Φ > 1) indicates weak H-bonds^54^. Cao et al. sought to derive Φ from the ratio of k_exp,HDX_ / k_exp,DHX_ and calculated H-bond strength ΔG from Φ, using an empirical relationship^42,55^. Although the profile of exchange rate constants in Cao et al. qualitatively matches the one found by us, the obtained ΔG values suggested strong H-bonds within TM-N while exceptionally weak H-bonds were reported at T43, V44, and T48 of TM-C. As this is at odds with the ΔG values reported here (Fig. 1c), we investigated the kinetic isotope effect in 80% TFE. First, we compared global DHX and HDX kinetics of WT and I45T, as obtained from MS measurements. DHX and HDX kinetics are superimposable after scaling the latter by a factor of ~0.2 (Supplementary Fig. S5a,b). The superimposability is consistent with similar k_exp,HDX_ / k_exp,DHX_ ratios for different populations of amides while the factor of 0.2 is reminiscent of the known 5-fold acceleration of the chemical DHX over HDX rate constants^56^. Second, we compared site-specific DHX and HDX rate constants. Since attempts to record HDX using ETD-MS were hampered by significant peak broadening, possibly resulting from hydrogen scrambling during gas phase fragmentation (data not shown), we recorded HDX and DHX by NMR spectroscopy (Supplementary Fig. S5c). As a result, the k_exp,HDX_ / k_exp,DHX_ ratios tended to be ≤1 from V40 through L52 (Supplementary Fig. S5d); ratios at all other positions were characterized by large errors since the associated quickly exchanging amides were difficult to capture at the timescale of the experiment (Supplementary Fig. S6 and S7). The potential sources of the discrepancies between our k_exp,HDX_ / k_exp,DHX_ ratios and the values reported by Cao et al.^42^ and the interpretation of these ratios are discussed below.

In summary, our results reveal a heterogeneous distribution of backbone H-bond strength *viz* helix flexibility across the APP TMD where a flexible TM-N is connected to a more rigid TM-C. The I45T mutation slightly stabilizes the helix upstream of the mutation site.

### I45T stabilizes the TM-C sub-helix in solution and in a POPC bilayer

To gain further mechanistic insights into the impact of the I45T mutation on the structural and dynamic properties of the APP TMD, we performed all-atom MD simulations in the μs range. A26-55 was simulated in 80% TFE and a POPC bilayer. No impact of I45T on the TMD’s orientation in the POPC bilayer was found (Supplementary Fig. S8). In investigating the helix-stabilizing network of intrahelical H-bonds, we first focused on occupancies, which represent the probability that an amide forms either an α- or a 3_10_ H-bond. In 80% TFE, the H-bond occupancies roughly follow the profile of exchange rates in DHX experiments, featuring a stable TM-C and a more flexible TM-N (Fig. 2a). Both sub-helices are connected by a region of low occupancy, spanning the region from O(L34) to NH(I41) and containing the G_37_G_38_ motif. As weak H-bonds enable structural deformations of the helix, this region of low occupancy triggers bending of the TMD over the G_37_G_38_ hinge^27,29,34^. In our simulations, the I45T mutation stabilized the amide H-bonds at the mutation site. Concomitantly, we found a slight destabilization of the H-bond emanating from I47 but unaffected stability at the ε-sites. The observed alterations of H-bond occupancies in the I45T mutant are related to an increased contribution of α H-bonds, while 3_10_ H-bonds contribute less than in WT (Supplementary Fig. S9a,b). Similar, albeit less pronounced, alterations of H-bond occupancies by I45T were detected in POPC (Fig. 2a and Supplementary Fig. S9a,b).

**Figure 2.**
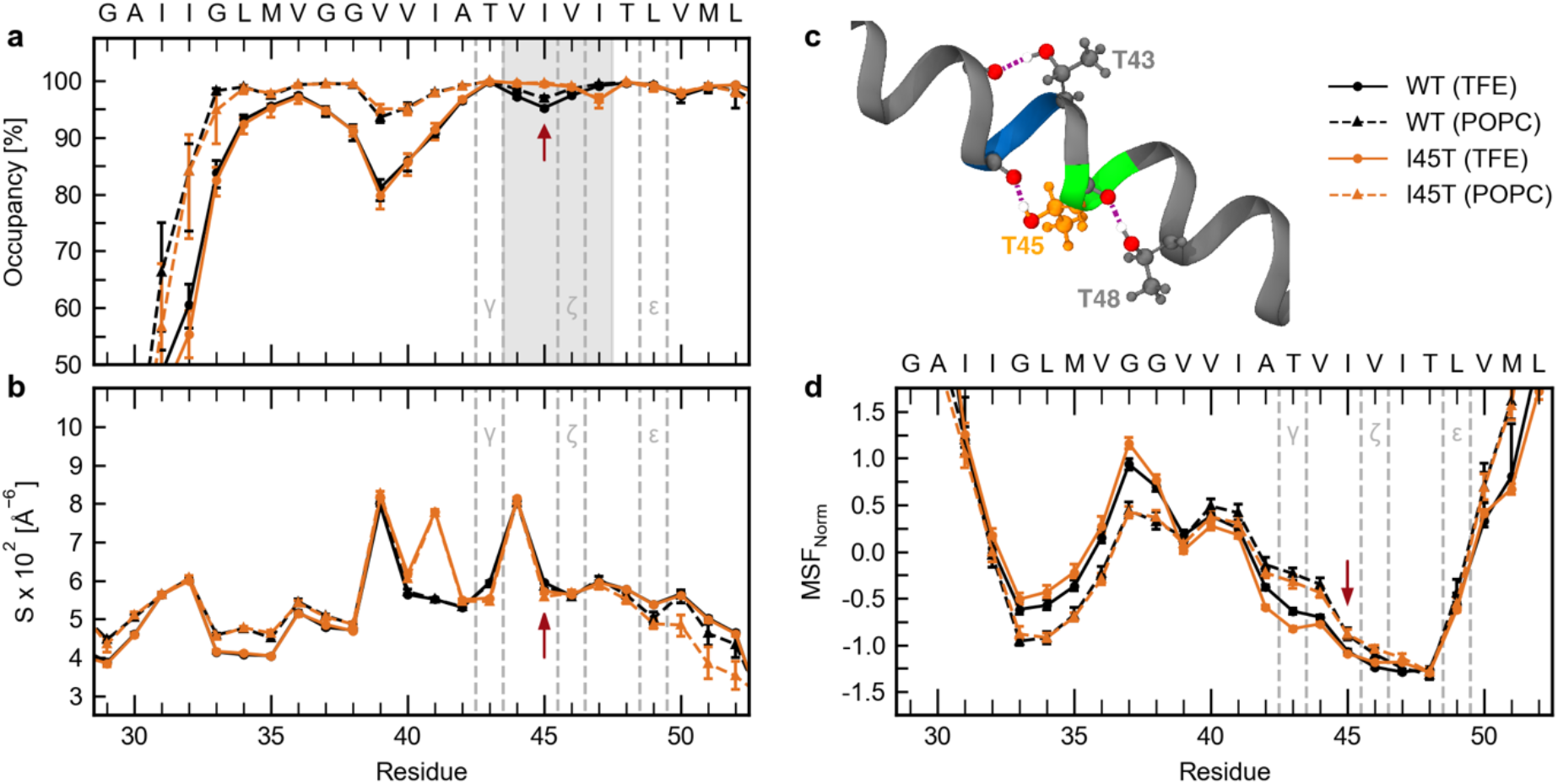
Local dynamics in 80% TFE and a POPC bilayer from MD simulations. (**a**) Occupancies of intramolecular H-bonds between the amide hydrogen at position i and carbonyl oxygens at position i-3 (3_10_ helix) or i-4 (α helix). Residues covered by the grey area showed significant differences between WT and I45T and were used as input for FMA. (**b**) Packing scores S_i_ along the TMD. (**c**) Visual representation of threonine back-bonding in the I45T mutant. Red spheres represent oxygen atoms and white spheres indicate hydrogen atoms involved in H-bonding. Green colour indicates ζ-cleavage sites while blue indicates γ-cleavage sites. Dashed violet lines represent H-bonds. (**d**) Mean-squared fluctuations normalized to zero mean and unit variance. Red arrows point to the I45T mutation site. Values represent means with 95% confidence intervals from bootstrap resampling.

The stabilities of amide H-bonds are influenced by side-chain interactions^57^. Therefore, we quantified the packing of side-chain heavy atoms around each backbone carbonyl oxygen by packing scores (Si). We found that a weakly packed TM-N was connected to a more tightly packed TM-C (Fig. 2b). Packing scores peak at V39 and V44, which is related to the back-bonding of T43 and T48, whose hydroxyl groups form H-bonds with the backbone carbonyl oxygens at the respective (i-4) positions (occupancy > 98%). As I45T harbours an additional Thr, a further peak at I41 reflects the additional H-bond between the T45 side chain and the I41 main chain. Packing profiles in POPC closely mirror the profiles found in 80% TFE. Together, this results in a highly coupled H-bond network around the ζ and γ cleavage sites as visualized in Fig. 2c.

Normalized mean-squared fluctuations (MSF) confirm above average flexibility at V_36_-V_40_ (Fig. 2d), as well as below average fluctuations in TM-N and TM-C. In particular, fluctuations at the ε-sites are lower than fluctuations of TM-N. Compared to 80% TFE, the POPC bilayer reduced fluctuations of TM-N and increased fluctuations of TM-C. Both in solution and in the bilayer, the I45T mutation decreased MSF from A42 through T45, but not at ε-sites. This finding is supported by indistinguishable distributions of rise-per residue values of Cα-atoms that flank the respective ε-cleavage sites (Supplementary Fig. S9c,d).

To summarize, an additional H-bond between the T45 side chain and the backbone carbonyl oxygen of I41 contributes to a tightly coupled H-bond network around the ζ- and γ-cleavage sites. This is related to an increased stability of intrahelical H-bonds at the mutation site and at positions upstream.

### I45T Alters Global Shape Fluctuations and Relocates Hinges in TM-C

Our investigation of intrahelical H-bonds revealed that the I45T mutation alters the relative importance of α and 3_10_ H-bonds (Supplementary Fig. S9a,b). Similar switching between α and 3_10_ H-bonds was attributed to shape deformations of TMD helices without disruption^58,59^. Therefore, we characterized shape deformations of the C99 TMD helix by analysing bending (θ) and swivel (Φ) angles between two helical segments. The first segment (I31-M35) centres around G33 in TM-N, which is in the middle of the G29xxxG33xxxG37 motif, potentially being involved in the binding of C99 to presenilin^33,34,60,61^. The second segment is located in TM-C (I_47_-M_51_) and harbours the ε-cleavage sites. Therefore, θ and Φ angles describe the positioning of the cleavage sites relative to a putative binding motif in TM-N. In agreement with previous studies^26,27,29,34^, the A26-55 WT helix bends anisotropically over the G_37_G_38_ hinge in 80% TFE (Fig. 3a,b). The I45T mutation leads to slightly increased bending angles and shifts the direction of bending by ~20° while reducing variations in bending direction (Fig. 3a-c). These differences between WT and I45T were reproduced in POPC, although the extent of bending was somewhat restricted in the membrane (Fig. 3c).

**Figure 3.**
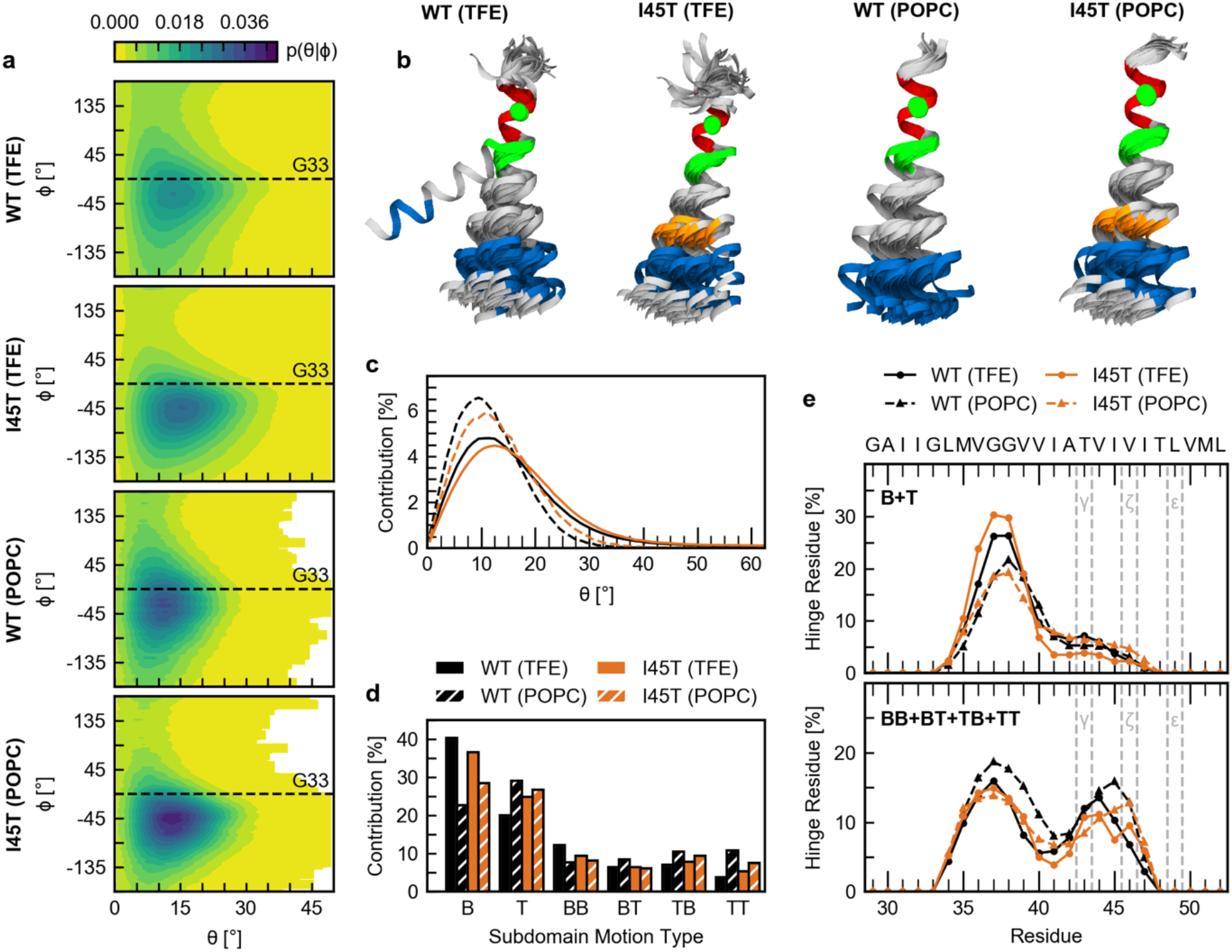
Collective TM-helix motions. (**a**) Bending (θ) and swivel (Φ) angles, which characterize the orientation of the ε-sites-carrying TM-C helix segment I47-M51 in relation to the TM-N segment I31-M35. (**b**) Representative conformations (overlaid at I31-M35) as determined by K-means clustering of θ and Φ angles (blue, TM-C segment I47-M51; red, TM-N segment I31-M35; orange, I45T mutation; green, G37 and G38; green sphere, C_α_ atom of G33). (**c**) Probability distributions of bending angles. (**d**) Probability of hinge motion types. Motions coordinated by a single hinge are referred to as type B and T, respectively. Motions around a pair of hinges are characterized by combinations of bending and twisting (BB, BT, TB, TT). (**e**) Probability by which a residue is identified as a hinge site in a single (B+T) and double-hinge (BB+BT+TB+TT) motion.

Our analysis of θ and Φ angles provides information about the orientation of the ε-cleavage sites in TM-C relative to TM-N. However, it lacks detailed information on the relative contributions of bending and twisting movements as well as the location of the dynamic regions, which coordinate them. The latter ones are commonly referred to as flexible hinges and are characterized as regions of low stability, flanked by two quasi-rigid segments. In order to detect the occurrence of hinges in the A26-55 TMD and to determine their locations as well as the type of motion they coordinate, we used the DynDom program^62^. Our analysis confirms a second hinge located in TM-C around T_43_-I_45_ in addition to the known G_37_G_38_ hinge (Fig. 3e).^29^ Each of these hinges can act as a flexible joint by itself (single-hinge bending (B) and twisting (T) motions) or cooperatively with each other (double-hinge motions of type BB, BT, TB, and TT). In 80% TFE, bending controlled by a single hinge (type B) is the dominant subdomain motion type (~ 40%) in the WT TMD, followed by a lower amount (~ 20%) of twisting (type T) (Fig. 3d). A similar preference for bending was noticed for the motions around the coupled pairs of hinges, where motion type BB, BT and TB contributed ~ 20% to overall backbone flexibility, while type TT motion contributed only ~ 5%. I45T increased the contribution of twisting by ~ 5%, while reducing bending by the same amount. The preference for twisting motions in POPC as well as the higher contribution of lower-amplitude double-hinge motions at the expense of large-scale single-hinge bending (B) is consistent with restricted conformational freedom in the membrane.

In addition to shifting the contributions of different types of motion to the overall backbone dynamics, we also observed an impact of I45T on the propensity by which each residue contributes to hinge flexibility (Fig. 3e and Supplementary Fig. S10). Single-hinge bending (B) and twisting (T) are mainly coordinated by residues V_36_-V_39_ in WT and I45T. However, the I45T mutation reduces the contribution of T_43_-I_45_ to hinge flexibility and slightly shifts propensities towards the C-terminus. In the POPC bilayer, hinge propensities around V_36_-V_39_ are reduced, while propensities around T_43_-I_45_ are slightly increased.

### H-Bond Dynamics Determines Motions Controlled by the Hinge at the ζ-Sites

Our analysis of hinges revealed that the differences between WT and I45T were mainly located around T_43_-I_45_ (Fig. 3e). This is in good agreement with observed alterations in H-bond stabilities (Fig. 2a). Therefore, we rationalize that alterations in H-bond open/close dynamics correlate with changes in hinge localization and the type of coordinated motion (Fig. 3d,e). As both are affected by I45T, motions controlled by such hinges were assumed to be of functional relevance for ε-site cleavage. In order to identify those functional motions (FM), we used a partial least square (PLS) model^63^ to find the motion which is maximally correlated with the collective open and close dynamics of H-bonds emanating from V_44_-I_47_. For WT, the FM in both environments is mainly determined by fluctuations of residues V_40_-I_45_. For the I45T mutant, residues T_43_-T_45_ in 80% TFE and residues A_42_-I_47_ in POPC mainly determine the FM. Thus, the ensemble of residues which collectively participate in the hinge motions is shifted towards the C-terminus in I45T (Fig. 4a), which is consistent with the shift of hinge sites. The DynDom program characterizes all FMs as motions with >50% bending. Hinge residues which coordinate this bending show the highest contribution to these FMs (green stars in Fig. 4a). The overlap between the FMs, which characterizes the similarity between two motions, reveals a significant difference between WT and I45T. FMs of either TMD in 80% TFE and POPC are similar (Fig. 4a, right panel). To visualize each peptide’s FM, conformations are interpolated along the FM (Fig. 4b,c). The FM of the WT helix in 80% TFE and in POPC consists of a bending motion, promoting a pronounced upward movement of the ε-sites (Fig. 4b). Surprisingly, such upward movement is not present in the I45T mutant. Top views on the interpolated conformations (Fig. 4c) exhibit only minor differences in the bending direction between WT and I45T in 80% TFE and in the POPC bilayer.

**Figure 4.**
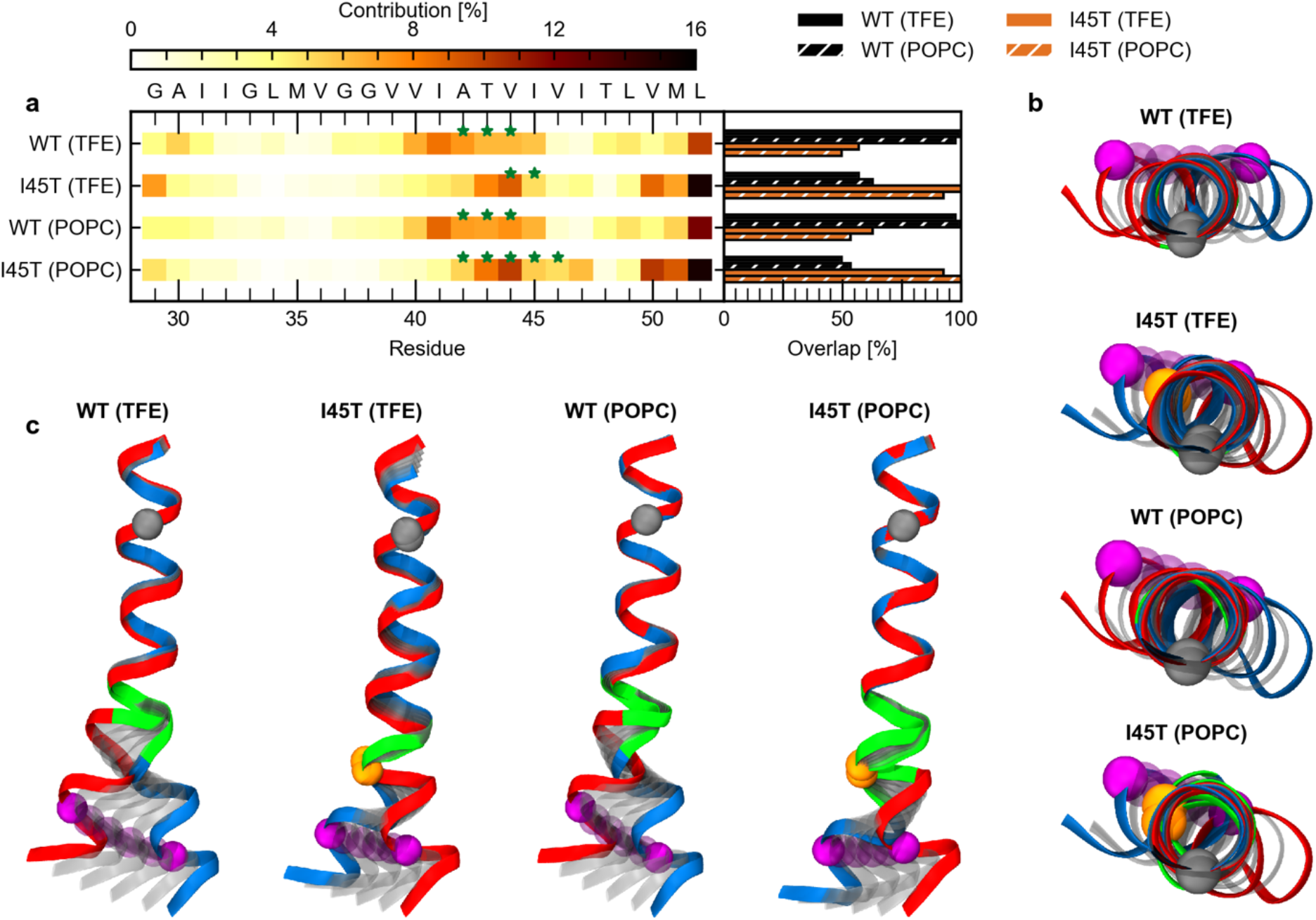
Functional model analysis. (**a**) Contribution of residues to the motion maximally correlated with occupancy variations of intrahelical amide H-bonds spanning residues V44 - I47. Green stars indicate residues that act as hinges. Overlap (inner product) between the individual ewMCM vectors is shown in the right panel. (**b**) Visualization of conformations as generated by interpolation along the ewMCM between maximum (red) and minimum (blue) extents. Structures were overlaid onto residues I31 – M35. Grey spheres represent the C_α_ atom of G33, orange spheres the C_α_ atom of T45 and purple spheres the C_α_ atom of L49. Residues classified as hinge residues (see **a**) are highlighted in green. (**c**) Top view on visualizations of the ewMCM as shown in **b**.

To summarize, observed alterations in H-bond dynamics by the I45T FAD mutation correlate with altered motion type and localization of hinge bending in TM-C. This motion induces an upward bending movement of the ε cleavage site in WT which is not present in the I45T mutant.

## Discussion

FAD mutations in the TM-C of the C99 TMD^19^ mainly lead to changes in the total efficiency of ε-cleavage, the preference for ε48 *vs*. ε49 sites, and the processivity of cleavage, thus altering Aβ ratios^15,20,36^. Neither the S’ pocket model of substrate binding^23,33^ nor the stability of the helix at the cleavage sites^28,31,33^, its global shape fluctuations^26^, or the location of dynamic elements (e.g., hinges) in the TMD^29^ offer a consistent explanation for these impacts of FAD mutations. The I45T mutation is a particular case as it strongly reduces ε-cleavage efficiency^20,21^ but does not significantly shift ε-site preference^23^. At the same time, I45T increases the Aβ42/Aβ40 ratio^20,21,23^ which indicates pathway switching after initial ε-cleavage^23^ and contradicts the S’ pocket model of substrate binding^23,33^, as the T45 side-chain should fit even better in the small S2’ pocket of the ε49-ζ46-γ43 pathway.

Here, were investigated the impact of the I45T FAD mutation on the local and global backbone dynamics of the C99 TMD. Our amide H-bond stability profiles indicate that the weakly packed TM-N of C99 is more flexible than the more tightly packed TM-C, which harbours the cleavage sites. Normalized MSF profiles confirm below-average fluctuations around the ε cleavage sites. Moreover, in addition to the known G_37_G_38_ hinge^34^, our simulations confirm the presence of a previously described second flexible hinge located in TM-C^29^. The I45T FAD mutation alters H-bond stability around and upstream of the mutation site and, thereby, relocates and stiffens the hinge in TM-C^26,29^.

In general, our results obtained *in silico* are in good agreement with our experimental work. Differences between them are mainly seen in TM-N. Since this part is very flexible, differences may originate from: (1) difficulties to capture very rapidly exchanging amides associated with uncertainties in the quantitative evaluation of the experimental data, (2) a less structured TM-N in the A28-55 peptides (relative to A26-55 peptides) used in the ETD-MS experiments, and/or (3) incomplete sampling of unfolded or slightly disordered regions in the simulations. Compared to our ETD-MS measurements, NMR spectroscopy suggests slower amide exchange within TM-N and somewhat faster exchange within parts of TM-C. A potential source of this discrepancy may be a less structured TM-N in the A28-55 peptides used in the ETD-MS experiments. A monomer/dimer equilibrium as the further source can be excluded because concentration effects were not discernible in the NMR experiments between 50 and 1000 μM (data not shown).

The higher flexibility of TM-N relative to TM-C is in conflict with recently reported H-bond strengths for the C99 TMD in LMPG micelles^42^. There, D/H fractionation factors Φ were derived from k_exp,HDX_ / k_exp,DHX_ ratios. Generally, the Φ value reports the isotopic preference in an H-bond, where the equilibrium enrichment of deuterium at the amide indicates weak H-bonds (Φ > 1). Cao et al. reported Φ < 1 in TM-N and Φ > 1 for some residues in TM-C, concluding that the C99 TMD has strong H-bonds in TM-N and weak H-bond for some residues (T43, V44, and T48) in TM-C^42^. Here, we show that k_exp,HDX_ / k_exp,DHX_ ratios determined in isotropic solution are close to 0.2 in overall MS-based amide exchange as well as in those site-specific NMR measurements where experimental error is sufficiently low. Indeed, Φ derived from the k_exp,HDX_ / k_exp,DHX_ ratios by Cao et al^42^ may not reflect the stability of an intrahelical amide-to-carbonyl H-bond. Rather, we propose that they reflect the stability of an amide-to-solvent H-bond which is determined by the chemical exchange rate constants k_ch_, as detailed in the Supplementary Discussion. This is supported by measurements of k_exp,HDX_ / k_exp,DHX_ ratios by Cao et al. and ourselves, many of which were close to the average k_ch,HDX_ / k_ch,DHX_ ratio of 0.2^56^. The definition of H-bond strength by Cao et al. thus appears to include amide H-bonding to the solvent. Concerning reported weak H-bonds for residues T43, V44, T48 in TM-C, it has to be noted, that conspicuous features of amide hydrogen D/H fractionation factors of threonine residues have been reported previously^54,64^. It was shown that electrostatic stabilization of cooperative H-bonds emanating from the amide and the side chain of a threonine dramatically decreases equilibrium fractionation factors. The dielectric environment influences the stiffness of amide vibrational modes and pKa differences between donor and acceptor and thus, the chemical rate constants, in an isotope-specific way. The effect will be more pronounced in the low-dielectric interior of the micelle (DK 4-10^65^) as compared to 80% TFE (DK approx. 35^66^). Therefore, the weak amide-to-solvent H-bonds reported for T43, V44 and T48 following the procedure given in Cao et al. are compatible with strong intrahelical H-bonds and back-bonding from threonine side chains.

What is the reason for the increased H-Bond stability in the TM-C of the I45T mutation relative to WT? Our MD simulations revealed an additional H-bond between the hydroxyl group of the T45 side chain and the backbone carbonyl oxygen of I41. Similar back-bonding interactions at T43 and T48^26,29^ had previously been shown to influence the extent and direction of the helix bending of the C99 TMD^26^. Mutating T43 or T48 alters ε-cleavage efficiency as well as Aβ42/Aβ40 ratios by shifting ε-site preference^15,20,21,36^. By introducing an additional H-bond, the I45T mutation strengthens the network of H-bonds in TM-C.

Bending fluctuations in the APP TMD are mainly controlled by a hinge located around G_37_G_38_^26,29,34^ which determines the orientation of TM-C, harbouring the cleavage sites, relative to TM-N, harbouring putative binding motifs^26,27,29^. The I45T mutation caused only minor variations of these fluctuations, which does not explain the severely reduced ε-cleavage efficiency^20,21^. Recently, we identified an additional flexible region around T_43_-I_45_, which can act as an additional flexible joint, controlling ε-site mobility relative to the centre of the helix^29^. Here, we found that this hinge controls the backbone motion that correlates maximally with the open/close dynamics of those H-bonds affected by the I45T mutation. Crucially, an upward movement of the ε-sites could be observed in the WT but was absent in I45T. The lack of this upward movement might explain the reduced ε-cleavage efficiency, as it may adjust the height of the ε-cleavage sites relative to the catalytic aspartates of presenilin.

The additional H-bond contributed by the T45 side chain might also explain the pathway switching observed in the I45T mutant^20,23^. As this side chain stabilizes the helix around the ζ- and γ-cleavage sites, back-bonding might inhibit unfolding within the catalytic cleft of presenilin and thus limit its access to the ζ-46 scissile bond. As a result, presenilin may switch from the ε49-ζ46-γ43 pathway to the ζ45-γ42 pathway which explains the almost complete lack of Aβ43 ^20^. We note that this is rather speculative, as the dynamics of the Aβ49 cleavage product in the catalytic site of γ-secretase is currently unknown and subject to future investigations. However, this model is supported by experimental results, which showed increased stability at the cleavage site to decrease cleavage efficiency in γ-secretase^33^ and even inhibit cleavage in rhomboids^67^.

In summary, the I45T mutation strengthens the H-bond network in TM-C and changes the position and dynamics of a flexible hinge that is located one turn upstream of the ε-sites. The combined effect may alter the access of both ε-sites by the catalytic Asp residues of the enzyme, thus providing a rationale for reduced ε-cleavage efficiency without affecting ε-site preference.

## Methods

### Peptide Sequences

In this study, we used short model peptides which consisted of either residues 26-55 of C99 (A26-55) or residues 28-55 of C99 (A28-55) with an additional KK tag at the terminus (Table 1). The A26-55 was used for overall exchange and NMR experiments as well as MD simulations. For ETD measurements, we had to use A28-55 in order to achieve proper fragmentation. Similar C99 TMD peptides were shown to be good substrates for γ-secretase^31,43^. For more native conditions, we blocked both terminal ends by acetylation (N-Ter) and amidation (C-Ter).

**Table 1.** Sequences of peptides investigated in this study

| Peptide | Mutation | Sequence |
| --- | --- | --- |
| A26-55 | WT | Ac-SNKGAIIGLMVGGVVIATVIVITLVMLKKK-NH<sub>2</sub> |
|  | I45T | Ac-SNKGAIIGLMVGLVVIATV**T**VITLVMLKKK-NH<sub>2</sub> |
| A28-55 | WT | Ac-**KKK**GAIIGLMVGGVVIATVIVITLVMLKKK-NH<sub>2</sub> |
|  | I45T | Ac-**KKK**GAIIGLMVGLVVIATV**T**VITLVMLKKK-NH<sub>2</sub> |

### Peptide Synthesis

Peptides were synthesized by Fmoc chemistry by PSL, Heidelberg, Germany and purified to >90% purity as judged by mass spectrometry. All other chemicals were purchased from Sigma-Aldrich Co. (St. Louis, Missouri, USA).

### Deuterium-Hydrogen Exchange by MS/MS

All mass spectrometric experiments were performed on a Synapt G2 HDMS (Waters Co., Milford, MA). Samples were injected from a 100 μL Hamilton gas-tight syringe via a Harvard Apparatus 11 Plus with a flow rate of 5 μL/min. Spectra were acquired in a positive-ion mode with one scan per second and a 0.1 s interscan time.

Solutions of deuterated peptide (100 μM in 80% (v/v) d1-trifluoroethanol (d1-TFE) in 2 mM ND4-acetate) were diluted 1:20 with protonated solvent (80% (v/v) TFE in 2 mM NH4-acetate, pH 5.0) to a final peptide concentration of 5 μM and incubated at a temperature of 20.0°C in a thermal cycler (Eppendorf, Germany). Incubation times were 0, 1, 2, 5, 10, 20, 30, 40, 50 min, and 1, 2, 3, 4, 6, 8, 12, 24, 48, 72 h. Exchange reactions were quenched by cooling the samples on ice and lowering the pH to 2.5 by adding 0.5% (v/v) formic acid. Mass/charge ratios were recorded and evaluated as previously described^48,49^, including a correction for the dilution factor. For electron transfer dissociation (ETD), we used 1,4-dicyanobenzene as a reagent on 5+ charged peptides, preselected via MS/MS. Fragmentation of peptides was performed as described in Stelzer et al.^35^. Briefly, ETD MS/MS scans were accumulated over a 10 min scan time, smoothed (Savitzky-Golay, 2 x 4 channels), and centred (80% centroid top, heights, 3 channels). ETD-measurements were performed after 13 different incubation periods (from 1 min to 3 d) where exchange took place at pH 5.0. Shorter (0.1 min, 0.5 min) and longer (5 d, 7 d) incubation periods were simulated by lowering the pH to 4.0 or elevating pH to 6.45, respectively, using matched periods. The differences to pH 5.0 were considered when calculating the corresponding rate constants. We note that base-catalysed exchange is responsible for at least 95% of the total deuteron exchange at ≥pH 4.0. The resulting ETD c and z fragment spectra were evaluated using a semi-automated procedure (ETD FRAGMENT ANALYZER module of MassMap_2017-11-16_LDK Software, MassMap GmbH & Co. KG, Freising). The free energies ΔG required for H-bond opening were calculated from k_exp,DHX_ and k_ch_ based on equation (1) based on Linderstrøm-Lang theory, assuming EX2 conditions and a predominantly folded state^52^.

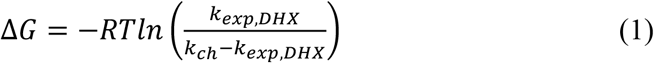

where k_ch_ represents the sequence-specific chemical rate constants that were calculated using the program SPHERE (http://landing.foxchase.org/research/labs/roder/sphere/) (under the set conditions: D-to-H-exchange, reduced Cys, pH =5.0, T = 20.0°C).

It should be noted, that the ΔG values obtained with this procedure are an upper estimate of the true values since (i) the molarity of water in 80% (v/v) TFE solvent is only 20% of the bulk molarity used for the determination of the reference chemical exchange rates kch, and (ii) the hydration of residues in the hydrophobic core of a TMD is possibly reduced relative to bulk. Both factors likely reduced the chemical exchange rate in our experiments. In addition, TFE might have an impact on the autoionization constant of water and the chemical exchange rate constants^35^. A detailed outline of the method is published elsewhere^53^.

Due to the 5% of deuterated solution in the DHX-ETD assay, we fitted the data by equation (2) to calculate k_exp,DHX_.

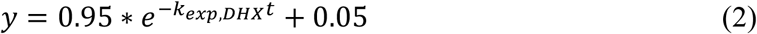

The extent of hydrogen scrambling could not be calculated with the ammonia loss method^68^ due to the blocked N-termini. However, previous experiments with similar peptides showed scrambling to be negligible under our conditions^59^. The absence of significant hydrogen scrambling is also indicated by the successful reconstruction of global exchange kinetics from the ETD data (Supplementary Fig. S3d,e).

### Molecular Dynamics Simulations

We performed molecular dynamics (MD) simulation of the A26-55 WT peptide and its I45T mutant (Table 1). Because no conformations were available for the investigated peptides, we developed a stochastic sampling protocol to generate a set of initial starting conformations (see Supplementary Methods).

For TFE/water, we performed a total of 78 simulations of 200 ns length using cluster centroids determined by affinity propagation clustering^69^ as input conformations. Settings, as described in Pester et al.^27^, were used for each simulation. In brief, each conformation was placed in a rectangular solvent box containing 80% TFE and 20% TIP3 (v/v). Equilibration was carried out in multiple steps by reducing harmonic restraints over a total of 1.2 ns. Production runs were performed in an NPT ensemble (T = 293 K, p = 0.1 MPa) using NAMD 2.11^70^ and the CHARMM36 force field^71^. An aggregated simulation time of 15.6 μs was collected for each peptide. The last 150 ns of each simulation were subjected to analysis, leading to an effective aggregated analysis over 11.7 μs for each peptide. Frames were recorded every 10 ps.

For simulations in POPC, a centroid conformation as obtained by hierarchical clustering (see Supplementary Methods) was placed in a POPC bilayer, consisting of 128 POPC lipids, using protocols as provided by CHARMM-GUI^72^. Simulations of 2.5 μs (T=303.15 K, p = 0.1 MPa) were performed, using NAMD 2.12^70^, the CHARMM36 force field^71^ and settings as provided by CHARMM-GUI. Frames were recorded every 10 ps. Only the last 1.5 μs of each trajectory were subjected to analysis.

### Analysis of MD Simulations

To validate the MD simulations, DHX kinetics were calculated from the MD simulations as described in Högel et. al.^59^. In order to account for non-deuterated residues in the experiment, the fit function was modified as shown in equation (3).

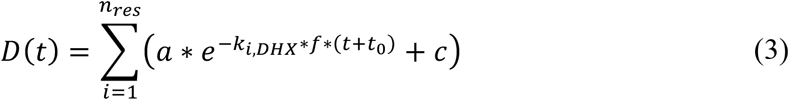

To account for 5% of non-deuterated peptide in the experiment, the amplitude (a) was set to 0.95 and a baseline (c) of 0.05 was added to the equation. In addition to the original protocol, we added a second fitting parameter t0 which accounts for time delays in the experiment. The quality of the MD-derived prediction of exchange kinetics was assessed by the normalized mean-squared deviation (χ^2^) of the averaged D(t) values with respect to the experimental averages.

Occupancies of closed H-bonds were computed for the types α, 3_10_ or helix (α or 3_10_ closed). Thereby, an H-bond was considered to be closed if the *O* ··· *H* distance was < 0.26 nm and the *O* ··· *H* – *N* angle was in the range 180° ± 60°.

Packing scores S_i_ measure the contacts of the carbonyl oxygen of residue i to all other atoms j and were computed as described in Götz & Scharnagl^29^.

Backbone mean-squared fluctuations (MSF, Cα atoms of residues G29-L52) were calculated for non-overlapping blocks of 30 ns. The block mean structure was calculated iteratively^73^. Normalization was done as described in Götz & Scharnagl^29^.

To access collective large-scale motions of the helix backbone, the bending Θ and swivel Φ angles between a helical segment in TM-N and TM-C, respectively, were computed as in Götz & Scharnagl^29^. The segment in TM-N covered residue I31-M35, while the segment in TM-C covered residues I47-M51.

Hinge-bending and twisting motions in the TM-helix were analysed by the Dyndom program^62^ as described in Götz & Scharnagl^29^. In contrast to the original protocol, snapshots were analysed every 100 ps.

Functional mode analysis was performed by the PLS-FMA program which uses a partial least-squares (PLS) model and was kindly provided by Bert de Groot^63^. Helix H-bond occupancies (α or 3_10_ closed) for residues V44-I47 were summed up and their time series were used as a functional order parameter for the model. Heavy backbone atoms of residues G29-L52 were used for the correlation analysis. The first half of the trajectory was used for model training, while the second half was used for cross-validation of the PLS-FMA model. The required number of PLS components was determined from the convergence of the Pearson correlation coefficient between data and model (R_m_) as a function of the number of components (Supplementary Fig. S11). Structural changes causing substantial variation in the order parameters were characterized by the ensemble-weighted, maximally correlated motions (ewMCM). For visualization, trajectories along the ewMCM vectors interpolating from low to high value of occupancies were used. Characterization of the ewMCM was done by DynDom, subjecting the conformations with the highest and lowest extent along ewMCM to analysis. Similarities of ewMCM vectors were quantified by the inner product of ewMCM vectors.

If not mentioned otherwise, all analyses used custom-built Python scripts based upon the MDtraj library^74^. Visuals were generated by VMD 1.9.2^75^.

### Statistical Evaluation of MD Results

Mean values and 95% confidence intervals were obtained by bias-corrected and accelerated bootstrap resampling^76^ of block averages. A block size of 30 ns was chosen to be > 2τ, with τ representing the autocorrelation function’s first zero passage time (Supplementary Fig. S12). Error propagation was performed by Monte-Carlo sampling. For resampling, values for each residue in a block were considered to be dependent and 10,000 samples were generated

### Data Availability

The datasets generated during and/or analysed during the current study are available from the corresponding author on reasonable request.

## Acknowledgments

This work was supported by the Deutsche Forschungsgemeinschaft (German Research Foundation) in the framework of the DFG research unit FOR2290 through grants LA699/20-1, SCHA630/4-1 and LU835/12-1 as well as by the Center for Integrated Protein Science Munich (CIPS^M^). The ETD FRAGMENT ANALYZER module of MassMap was kindly provided by Manfred Wozny (MassMap GmbH & Co. KG, Freising). The lsNMR experiments in this work were funded by the Helmholtz Programme BioInterfaces in Technology and Medicine (BIFTM) of the Karlsruhe Institute of Technology (KIT) and the DFG instrumental facility Pro^2^NMR. Computing resources were provided by the Leibniz Supercomputing Centre (LRZ) through grants ta511 (Linux Cluster) and pr92so (SuperMUC) and by the Gauss Centre for Supercomputing (GCS) through grants pr48ko (SuperMUC, LRZ). We want to thank Harald Steiner for his comments on the manuscript.

## Author Contributions

AG performed MD simulations, analysed the data and created the figures. PH performed mass spectrometry, analysed the data and contributed supplementary figures and tables with assistance from IH. MS performed NMR experiments, analysed the data and contributed supplementary figures. CS contributed theoretical thoughts about the isotope effect. DL, CS, CMG and BL designed and supervised the project. AG and PH wrote the manuscript with assistance from CS and DL. All authors have commented and given approval to the final version of the manuscript.

## Additional Information

**Supporting Information:** Supporting information contains additional discussion, methods, Tables S1-S2 and Figures S1-S12.

**Competing Interests:** The authors declare no competing financial interests.

## References

1. Haapasalo, A. & Kovacs, D. M. The many substrates of presenilin/γ-secretase. J. Alzheimers. Dis. 25, 3–28 (2011).

2. Beel, A. J. & Sanders, C. R. Substrate specificity of gamma-secretase and other intramembrane proteases. Cell. Mol. Life Sci. 65, 1311–34 (2008).

3. Haass, C., Kaether, C., Thinakaran, G. & Sisodia, S. Trafficking and Proteolytic Processing of APP. Cold Spring Harb. Perspect. Med. 2, a006270–a006270 (2012).

4. Kaether, C., Haass, C. & Steiner, H. Assembly, trafficking and function of γ-secretase. Neurodegener. Dis. 3, 275–283 (2006).

5. Lichtenthaler, S. F., Haass, C. & Steiner, H. Regulated intramembrane proteolysis - lessons from amyloid precursor protein processing. J. Neurochem. 117, 779–796 (2011).

6. Fukumori, A., Fluhrer, R., Steiner, H. & Haass, C. Three-Amino Acid Spacing of Presenilin Endoproteolysis Suggests a General Stepwise Cleavage of γ-Secretase-Mediated Intramembrane Proteolysis. J. Neurosci. 30, 7853–7862 (2010).

7. Matsumura, N. et al. γ-Secretase Associated with Lipid Rafts. J. Biol. Chem. 289, 5109–5121 (2014).

8. Olsson, F. et al. Characterization of intermediate steps in amyloid beta (Aβ) production under near-native conditions. J. Biol. Chem. 289, 1540–50 (2014).

9. Qi-Takahara, Y. et al. Longer Forms of Amyloid Protein: Implications for the Mechanism of Intramembrane Cleavage by -Secretase. J. Neurosci. 25, 436–445 (2005).

10. Quintero-Monzon, O. et al. Dissociation between the Processivity and Total Activity of γ-Secretase: Implications for the Mechanism of Alzheimer’s Disease-Causing Presenilin Mutations. Biochemistry 50, 9023–9035 (2011).

11. Takami, M. et al. γ-Secretase: Successive Tripeptide and Tetrapeptide Release from the Transmembrane Domain of β-Carboxyl Terminal Fragment. J. Neurosci. 29, 13042–52 (2009).

12. Saito, T., Matsuba, Y., Yamazaki, N., Hashimoto, S. & Saido, T. C. Calpain Activation in Alzheimer’s Model Mice Is an Artifact of APP and Presenilin Overexpression. J. Neurosci. 36, 9933–9936 (2016).

13. Sandebring, A., Welander, H., Winblad, B., Graff, C. & Tjernberg, L. O. The Pathogenic Aβ43 Is Enriched in Familial and Sporadic Alzheimer Disease. PLoS One 8, e55847 (2013).

14. Selkoe, D. J. & Hardy, J. The amyloid hypothesis of Alzheimer’s disease at 25 years. EMBO Mol. Med. 8, 595–608 (2016).

15. Dimitrov, M. et al. Alzheimer’s disease mutations in APP but not γ-secretase modulators affect epsilon-cleavage-dependent AICD production. Nat. Commun. 4, 2246 (2013).

16. Kakuda, N. et al. Equimolar Production of Amyloid β-Protein and Amyloid Precursor Protein Intracellular Domain from β-Carboxyl-terminal Fragment by γ-Secretase. J. Biol. Chem. 281, 14776–14786 (2006).

17. Page, R. C. et al. β-Amyloid Precursor Protein Mutants Respond to γ-Secretase Modulators. J. Biol. Chem. 285, 17798–17810 (2010).

18. Weggen, S. & Beher, D. Molecular consequences of amyloid precursor protein and presenilin mutations causing autosomal-dominant Alzheimer’s disease. Alzheimers. Res. Ther. 4, 9 (2012).

19. Alzforum. Mutations Database. (2018). Available at: https://www.alzforum.org/mutations. (Accessed: 9th January 2018)

20. Chávez-Gutiérrez, L. et al. The mechanism of γ-Secretase dysfunction in familial Alzheimer disease. EMBO J. 31, 2261–2274 (2012).

21. Xu, T.-H. et al. Alzheimer’s disease-associated mutations increase amyloid precursor protein resistance to γ-secretase cleavage and the Aβ42/Aβ40 ratio. Cell Discov. 2, 16026 (2016).

22. Sato, T. et al. Potential Link between Amyloid β-Protein 42 and C-terminal Fragment 49-99 of β-Amyloid Precursor Protein. J. Biol. Chem. 278, 24294–24301 (2003).

23. Bolduc, D. M., Montagna, D. R., Seghers, M. C., Wolfe, M. S. & Selkoe, D. J. The amyloid-beta forming tripeptide cleavage mechanism of γ-secretase. Elife 5, 1–4 (2016).

24. Dominguez, L., Foster, L., Straub, J. E. & Thirumalai, D. Impact of membrane lipid composition on the structure and stability of the transmembrane domain of amyloid precursor protein. Proc. Natl. Acad. Sci. 113, E5281–E5287 (2016).

25. Pester, O., Götz, A., Multhaup, G., Scharnagl, C. & Langosch, D. The Cleavage Domain of the Amyloid Precursor Protein Transmembrane Helix Does Not Exhibit Above-Average Backbone Dynamics. ChemBioChem 14, 1943–1948 (2013).

26. Scharnagl, C. et al. Side-Chain to Main-Chain Hydrogen Bonding Controls the Intrinsic Backbone Dynamics of the Amyloid Precursor Protein Transmembrane Helix. Biophys. J. 106, 1318–1326 (2014).

27. Pester, O. et al. The Backbone Dynamics of the Amyloid Precursor Protein Transmembrane Helix Provides a Rationale for the Sequential Cleavage Mechanism of γ-Secretase. J. Am. Chem. Soc. 135, 1317–1329 (2013).

28. Sato, T. et al. A helix-to-coil transition at the ε-cut site in the transmembrane dimer of the amyloid precursor protein is required for proteolysis. Proc. Natl. Acad. Sci. 106, 1421–1426 (2009).

29. Götz, A. & Scharnagl, C. Dissecting conformational changes in APP’s transmembrane domain linked to ε-efficiency in familial Alzheimer’s disease. PLoS One 13, e0200077 (2018).

30. Strisovsky, K. Why cells need intramembrane proteases - a mechanistic perspective. FEBS J. 283, 1837–1845 (2016).

31. Chen, W. et al. Familial Alzheimer’s mutations within APPTM increase Aβ42 production by enhancing accessibility of ε-cleavage site. Nat. Commun. 5, 3037 (2014).

32. Lu, J.-X., Yau, W.-M. & Tycko, R. Evidence from Solid-State NMR for Nonhelical Conformations in the Transmembrane Domain of the Amyloid Precursor Protein. Biophys. J. 100, 711–719 (2011).

33. Fernandez, M. A. et al. Transmembrane Substrate Determinants for γ-Secretase Processing of APP CTFβ. Biochemistry 55, 5675–5688 (2016).

34. Barrett, P. J. et al. The Amyloid Precursor Protein Has a Flexible Transmembrane Domain and Binds Cholesterol. Science (80-.). 336, 1168–1171 (2012).

35. Stelzer, W., Scharnagl, C., Leurs, U., Rand, K. D. & Langosch, D. The Impact of the ‘Austrian’ Mutation of the Amyloid Precursor Protein Transmembrane Helix is Communicated to the Hinge Region. ChemistrySelect 1, 4408–4412 (2016).

36. Oestereich, F. et al. Impact of Amyloid Precursor Protein Hydrophilic Transmembrane Residues on Amyloid-Beta Generation. Biochemistry 54, 2777–2784 (2015).

37. Ma, B. & Nussinov, R. Enzyme dynamics point to stepwise conformational selection in catalysis. Curr. Opin. Chem. Biol. 14, 652–9 (2010).

38. Agarwal, P. K., Doucet, N., Chennubhotla, C., Ramanathan, A. & Narayanan, C. in 273–297 (2016). doi:10.1016/bs.mie.2016.05.023

39. Miyashita, N., Straub, J. E. & Thirumalai, D. Structures of β-Amyloid Peptide 1–40, 1–42, and 1–55—the 672–726 Fragment of APP—in a Membrane Environment with Implications for Interactions with γ-Secretase. J. Am. Chem. Soc. 131, 17843–17852 (2009).

40. Beel, A. J. et al. Structural Studies of the Transmembrane C-Terminal Domain of the Amyloid Precursor Protein (APP): Does APP Function as a Cholesterol Sensor? † ‡. Biochemistry 47, 9428–9446 (2008).

41. Dominguez, L., Meredith, S. C., Straub, J. E. & Thirumalai, D. Transmembrane Fragment Structures of Amyloid Precursor Protein Depend on Membrane Surface Curvature. J. Am. Chem. Soc. 136, 854–857 (2014).

42. Cao, Z., Hutchison, J. M., Sanders, C. R. & Bowie, J. U. Backbone Hydrogen Bond Strengths Can Vary Widely in Transmembrane Helices. J. Am. Chem. Soc. 139, 10742–10749 (2017).

43. Yin, Y.I.et al. γ-Secretase Substrate Concentration Modulates the Aβ42/Aβ40 Ratio. J. Biol. Chem. 282, 23639–23644 (2007).

44. Sato, C., Morohashi, Y., Tomita, T. & Iwatsubo, T. Structure of the catalytic pore of gamma-secretase probed by the accessibility of substituted cysteines. J. Neurosci. 26, 12081–8 (2006).

45. Schutz, C. N. & Warshel, A. What are the dielectric ‘constants’ of proteins and how to validate electrostatic models? Proteins: Structure, Function and Genetics 44, 400–417 (2001).

46. Tolia, A., Chávez-Gutiérrez, L. & De Strooper, B. Contribution of Presenilin Transmembrane Domains 6 and 7 to a Water-containing Cavity in the γ-Secretase Complex. J. Biol. Chem. 281, 27633–27642 (2006).

47. Buck, M. Trifluoroethanol and colleagues: cosolvents come of age. Recent studies with peptides and proteins. Q. Rev. Biophys. 31, 297–355 (1998).

48. Stelzer, W., Poschner, B. C., Stalz, H., Heck, A. J. & Langosch, D. Sequence-specific conformational flexibility of SNARE transmembrane helices probed by hydrogen/deuterium exchange. Biophys J 95, 1326–1335 (2008).

49. Poschner, B. C., Quint, S., Hofmann, M. W. & Langosch, D. Sequence-specific conformational dynamics of model transmembrane domains determines their membrane fusogenic function. J Mol Biol 386, 733–741 (2009).

50. Xiao, H. Mapping protein energy landscapes with amide hydrogen exchange and mass spectrometry: I. A generalized model for a two-state protein and comparison with experiment. Protein Sci. 14, 543–557 (2005).

51. Konermann, L., Pan, J. & Liu, Y.-H. Hydrogen exchange mass spectrometry for studying protein structure and dynamics. Chem. Soc. Rev. 40, 1224–1234 (2011).

52. Skinner, J. J., Lim, W. K., Bédard, S., Black, B. E. & Englander, S. W. Protein dynamics viewed by hydrogen exchange. Protein Sci. 21, 996–1005 (2012).

53. Yücel, S. S. et al. Metastable XBP1u transmembrane domain mediates insertion into the ER membrane and intramembrane proteolysis by the signal peptide peptidase Sara. bioRxiv (2018). doi:10.1101/322107

54. Loh, S. N. & Markley, J. L. Hydrogen Bonding in Proteins As Studied by Amide Hydrogen D/H Fractionation Factors: Application to Staphylococcal Nuclease. Biochemistry 33, 1029–1036 (1994).

55. Cao, Z. & Bowie, J. U. An energetic scale for equilibrium H/D fractionation factors illuminates hydrogen bond free energies in proteins. Protein Sci. 23, 566–575 (2014).

56. Teilum, K., Kragelund, B. B. & Poulsen, F. M. in Protein Folding Handbook 634–672 (Wiley-VCH Verlag GmbH). doi:10.1002/9783527619498.ch18

57. Quint, S. et al. Residue-specific side-chain packing determines the backbone dynamics of transmembrane model helices. Biophys. J. 99, 2541–2549 (2010).

58. Cao, Z. & Bowie, J. U. Shifting hydrogen bonds may produce flexible transmembrane helices. Proc. Natl. Acad. Sci. 109, 8121–8126 (2012).

59. Högel, P. et al. Glycine Perturbs Local and Global Conformational Flexibility of a Transmembrane Helix. Biochemistry 57, 1326–1337 (2018).

60. Kornilova, A. Y., Bihel, F., Das, C. & Wolfe, M. S. The initial substrate-binding site of γ-secretase is located on presenilin near the active site. Proc. Natl. Acad. Sci. 102, 3230–3235 (2005).

61. Yan, Y., Xu, T.-H., Melcher, K. & Xu, H. E. Defining the minimum substrate and charge recognition model of gamma-secretase. Acta Pharmacol. Sin. 1–13 (2017). doi:10.1038/aps.2017.35

62. Hayward, S. & Lee, R. A. Improvements in the analysis of domain motions in proteins from conformational change: DynDom version 1.50. J. Mol. Graph. Model. 21, 181–183 (2002).

63. Krivobokova, T., Briones, R., Hub, J. S., Munk, A. & de Groot, B. L. Partial Least-Squares Functional Mode Analysis: Application to the Membrane Proteins AQP1, Aqy1, and CLC-ec1. Biophys. J. 103, 786–796 (2012).

64. Edison, A. S., Weinhold, F. & Markley, J. L. Theoretical Studies of Protium/Deuterium Fractionation Factors and Cooperative Hydrogen Bonding in Peptides. J. Am. Chem. Soc. 117, 9619–9624 (1995).

65. L’Heureux, G. P. & Fragata, M. Micropolarities of lipid bilayers and micelles. J. Colloid Interface Sci. 117, 513–522 (1987).

66. Gente, G. & La Mesa, C. Water-trifluoroethanol mixtures: Some physicochemical properties. J. Solution Chem. 29, 1159–1172 (2000).

67. Brown, M. C. et al. Unwinding of the Substrate Transmembrane Helix in Intramembrane Proteolysis. Biophys. J. 114, 1579–1589 (2018).

68. Rand, K. D., Zehl, M., Jensen, O. N. & Jorgensen, T. J. Loss of ammonia during electron-transfer dissociation of deuterated peptides as an inherent gauge of gas-phase hydrogen scrambling. Anal Chem 82, 9755–9762 (2010).

69. Frey, B. J. & Dueck, D. Clustering by passing messages between data points. Science 315, 972–976 (2007).

70. Phillips, J. C. et al. Scalable molecular dynamics with NAMD. J. Comput. Chem. 26, 1781–802 (2005).

71. Best, R. B. et al. Optimization of the additive CHARMM all-atom protein force field targeting improved sampling of the backbone φ, ψ and side-chain χ(1) and χ(2) dihedral angles. J. Chem. Theory Comput. 8, 3257–3273 (2012).

72. Lee, J. et al. CHARMM-GUI Input Generator for NAMD, GROMACS, AMBER, OpenMM, and CHARMM/OpenMM Simulations Using the CHARMM36 Additive Force Field. J. Chem. Theory Comput. 12, 405–413 (2016).

73. Romo, T. D. & Grossfield, A. Block Covariance Overlap Method and Convergence in Molecular Dynamics Simulation. J. Chem. Theory Comput. 7, 2464–2472 (2011).

74. McGibbon, R. T. et al. MDTraj: A Modern Open Library for the Analysis of Molecular Dynamics Trajectories. Biophys. J. 109, 1528–1532 (2015).

75. Humphrey, W., Dalke, A. & Schulten, K. VMD: Visual molecular dynamics. J. Mol. Graph. 14, 33–38 (1996).

76. DiCiccio, T. J. et al. Better Bootstrap Confidence Intervals. Stat. Sci. 11, 189–228 (1996).

